# Using citizen science to protect threatened amphibian populations in urban spaces: ecology, life history, and conservation of green toads and fire salamanders in Jerusalem and Haifa

**DOI:** 10.1101/2023.07.19.549780

**Authors:** Omer Darel, Olga Rybak, Asaf Ben Levy, Gabi Kolodny, Tamar Kis- Papo, Nirit Lavie Alon, Rotem Vidan, Oren Kolodny

**Affiliations:** Department of Ecology, Evolution and Behavior, The A. Silberman Institute of Life Sciences, The Hebrew University of Jerusalem, Jerusalem 9190401, Israel; The Society for the Protection of Nature in Israel; The Israeli Ministry of Education

## Abstract

1. The rapid urbanization processes occurring worldwide are amongst the main factors driving the current biodiversity crisis. In particular, a third of known amphibian species are directly threatened by urbanization. The negation of this threat will require conservation efforts aimed at sustaining viable amphibian populations within the urban landscape, which must be informed by a deep understanding of the way amphibian populations are affected by the unique risk factors of the urban environment.
2. To address this need for four populations of amphibians in Israel, we performed a capture-recapture analysis on two datasets. The larger of the two datasets is the result of a multi-year citizen science program focused on two *Salamandra infraimmaculata* populations within the city of Haifa, Israel. The second dataset is the result of one year of survey following a similar protocol that we performed on two *Bufotes variabilis* populations within the city of Jerusalem and at a nature reserve near it. Individuals of both species have unique and recognizable dorsal spot patterns, which allowed for noninvasive recapture identification.
3. The results of our analysis provide insights that can guide future conservation of the specific studied population, but our conclusions have wider implications, regarding both the ecology of the studied species and applied conservation science: using the salamander dataset, we developed a method of length-based age estimations for this species and found that the studied salamanders have a prolonged period of increased vulnerability throughout their first years of life, even after reaching sexual maturity.
4. Additionally, the shared conclusions from the two case studies indicate that the creation of fish-containing artificial water bodies in Mediterranean habitats can have detrimental impacts on the resident amphibian populations.
5. Synthesis and implications: The significance and extent of our results demonstrate the effectiveness of citizen science as a tool for research and conservation in the urban environment. Our findings call for the implementation of management practices that prioritize the protection of urban amphibians and their habitats. By identifying the vulnerability of amphibians during critical life stages and highlighting the negative impacts of fish-containing water bodies, our study contributes to the development of informed conservation policies with implications for urban planning, habitat management, and biodiversity conservation strategies.

## 1. Introduction

### 1.1 background and research goals

The world is urbanizing at an increasing rate)Desa, 2018; Seto et al., 2013(, a process that threatens ecosystems globally (Lin & Fuller, 2013). Amphibians, which are the most threatened land vertebrates (Cushman, 2006; Hamer & McDonnell, 2008; Stuart et al., 2004), are particularly sensitive to the environmental alterations driven by urbanization (Clevenot et al., 2018; Cushman, 2006; Hamer & McDonnell, 2008; Hamer & Parris, 2011; Schoch, 2009). Amphibians usually require both an aquatic and a terrestrial habitat to complete their life cycle, and their permeable skin and low mobility make them particularly vulnerable to the pollution, fragmentation, and habitat degradation which are characteristic of the urban landscape (Hamer & McDonnell, 2008). The ecosystems within the urban environment have traditionally been ignored by ecologists, but in recent years the interest in urban ecosystems has been increasing (Colléony & Shwartz, 2019; Lin & Fuller, 2013). This increase is partially due to the growing understanding of urban ecosystems’ vitality for mental health, and the importance of regular exposure to nature for maintaining positive views and sentiments towards nature, and therefore public support of, and participation in, conservation efforts (Colléony & Shwartz, 2019; Kay et al., 2022; Miller, 2005; Pyle, 2002, 2003). The abundance of studies resulting from this increased interest have enriched our knowledge and understanding of urban ecosystems (Beninde et al., 2015; Shwartz et al., 2013, 2014).

Several studies have shown that urban ecosystems frequently contain large biodiversity, and in particular, a comparable number of amphibian species to that found in rural ecosystems (Hamer & McDonnell, 2008; Hamer & Parris, 2011; Parris, 2006). However, their presence in urban ecosystems does not necessarily indicate that these species can sustain viable populations in there(Clevenot et al., 2018; Hamer & McDonnell, 2008). Specifically, the suitability of aquatic breeding sites such as stormwater reservoirs and garden ponds for amphibian reproduction have been studied, with mixed findings (Clevenot et al., 2018; Hamer & McDonnell, 2008). These studies raised the concern that urban water bodies may function as ecological traps: habitats, usually artificial or altered by human activity, which cannot support a certain species but are erroneously identified by that species as suitable, and by attracting individuals from suitable adjacent habitats, damage the local population (Clevenot et al., 2018; Demeyrier et al., 2016). Understanding the way urban environments can sustain viable, functioning populations of endangered amphibian species can be a vital tool in the mitigation of the threats posed to amphibians by the increasing rate of urbanization(Hamer & McDonnell, 2008; Oertli & Parris, 2019; Vignoli et al., 2009). To achieve this, it is crucial to extensively measure proxies of population viability, such as reproductive success, juvenile recruitment, and adult survival rate in urban amphibian populations (Clevenot et al., 2018; Hamer & McDonnell, 2008; Hamer & Parris, 2011; Parris, 2006). However, linking these attributes to the environmental conditions proves to be rather challenging; Urban breeding sites, such as stormwater reservoirs, can sustain a viable population in one setting, while functioning as an ecological trap in another, with no single dominant factor dictating the result, as demonstrated by a recent review by Clevenot et al., (2018) et al. Additionally, Clevenot’s study, like most studies of urban amphibians, was focused on North America and Europe, limiting the existing knowledge’s applicability to other parts of the world (Oertli & Parris, 2019).

The ecology of amphibians in the semiarid Mediterranean significantly differs from the well-studied temperate zones (Blank & Blaustein, 2012): Most of the breeding occurs in ephemeral ponds and small rock pools sustained by rainfall or small springs, both of which usually remain dry for most of the year (Degani & Kaplan, 1999; Ferreira & Beja, 2013); Unsurprisingly, the tadpoles of amphibians in the Mediterranean frequently possess phenotypic plasticity allowing them to adjust the timing of their metamorphosis to the unpredictable hydroperiods of their breeding sites (Goldberg et al., 2012); Additionally, the threat of desiccation posed by the harsh summers leads many amphibians to estivate in burrows or karstic cavities for months at a time (Degani & Warburg, 1978; Elron, 2007; Ferreira & Beja, 2013). The results of some studies indicate to some degree that the limiting factor for many of these populations is the availability of aquatic breeding sites (Ferreira & Beja, 2013; Vignoli et al., 2009).

There are eight known amphibian species in Israel, only one of which is not locally threatened (Dolev & Perevolotsky, 2004; Meiri et al., 2019); for most of these species Israel constitutes the southern edge of their distribution (Degani et al., 2019). Edge populations are generally under-studied (Cahill et al., 2013), and while these populations are often the most threatened by climate change(Cahill et al., 2013; Hampe & Petit, 2005), edge populations were speculated to be invaluable for the survival and evolution of species during climatic shifts(Hampe & Petit, 2005).

The projected population density and resulting urbanization rates in Israel are exceedingly high (Strategic Plan for Housing (Hebrew), 2017; OECD, 2022; Worldometers.info, 2022), and it is estimated that since the beginning of the 20th century 95% of the preexisting ephemeral water bodies in the country have been lost to urban and agricultural development, with the remaining aquatic habitats often being highly altered or degraded (Levin et al., 2009; Mendelssohn, 1985). The expected effects of climate change on the ephemeral water bodies in Israel are also abnormally severe, with an expected decrease in precipitation and intense warming leading to a rapid shortening of their hydroperiods (Givati & Rosenfeld, 2013).

We suggest that it is likely that alternative habitats within urban ecosystems will prove to be critical for the continued existence of Israel’s amphibians. Due to its location in a region that is expected to experience some of the most significant climatic shifts in the coming decades, it may serve as a possible indicator of the changes expected in other parts of the world (Givati & Rosenfeld, 2013). Information on the viability of urban amphibian populations in Israel is nevertheless lacking (Degani & Kaplan, 1999; Dolev & Perevolotsky, 2004; Oertli & Parris, 2019), and data on population sizes, breeding success, and adult survivorship is generally scarce for amphibians in Israel, and even more so in its urban environments (Segev et al., 2010).

To ensure that urban habitats’ potential as a tool in amphibian conservation is realized, we must develop our understanding of the factors determining their suitability for sustaining viable amphibian populations (Clevenot et al., 2018; Hamer & McDonnell, 2008; Hamer & Parris, 2011; Parris, 2006; Semlitsch, 2002). Roads, the introduction of predatory fish, and invasive amphibian competitors are some of the most consistent hazards recognized by previous studies, but have been shown to affect different amphibian species in varying, sometimes opposite ways (Clevenot et al., 2018; Hamer & McDonnell, 2008; Hamer & Parris, 2011; Kats & Ferrer, 2003). While most existing literature is focused on the community level, population-focused studies, as well as a better understanding of species-specific habitat utilization, movement patterns, and life history traits, are necessary to allow us to recognize the specific factors that jeopardize these species’ continued existence in the urban landscape (Hamer & McDonnell, 2008; Scheffers & Paszkowski, 2012). The study of movement patterns, population dynamics, and life history traits of the small, nocturnal, and sometimes cryptic amphibian species of the levant usually requires long-term and labor-intensive surveys, limiting the ability of many researchers in academia to provide this crucial information (Hamer & McDonnell, 2008; Parris, 2006; Scheffers & Paszkowski, 2012).

The use of citizen science in ecology has long been established as an effective tool for both research and education (Campbell et al., 1999; Palmer et al., 2021; Rae et al., 2019; Rowley et al., 2019; Sullivan et al., 2014; Westgate et al., 2015). In a rapidly urbanizing world, an increasing number of people are denied the opportunity to have regular contact with nature, a phenomenon dubbed the “extinction of experience” (EOE), with worrying implications for the public’s physical and mental health, as well as for their emotional involvement in conservation (Colléony et al., 2020; Colléony & Shwartz, 2019; Miller, 2005; Pyle, 2003). People who lack contact with, and knowledge of, amphibians in their vicinity, were shown to have a greater chance to have a negative disposition towards the amphibians in their environment, occasionally leading them to directly harm the amphibians they encounter (Pavol & Fančovičová, 2012; Riós-Orjuela et al., 2020; Sousa et al., 2016). The hands-on experience provided by participating in these citizen science projects was found to promote understanding and sympathy towards amphibians, among other wildlife (Sousa et al., 2016). In multiple cases, the participants of citizen science projects reported that their knowledge of the focal species, as well as their willingness to share this knowledge with other members of their community, or participate in conservation efforts of the focal species, have all increased after they participated in these programs (Peter et al., 2019).

In this paper we first present a case study of an intensive, multi-year citizen science project, which focused on two urban salamander populations in the city of Haifa, Israel. This project was conceived for the purpose of gathering as much information as possible on these populations to inform future conservation efforts. The shared efforts of dozens of local volunteers over six years allowed for locating and documenting large numbers of salamanders regularly, with multiple captures of the same individuals identified using their unique dorsal spot patterns. The extensive dataset collected in this project allowed us to perform a multi-year capture-recapture analysis on both populations, gathering insights on the population size, structure, and stability in the study sites, as well as deepening our understanding of the life history, movement patterns, and population ecology of these urban amphibians. Alongside this dataset, we analyzed a capture-recapture survey following a similar protocol on a smaller scale, which we conducted on two toad populations within and near the city of Jerusalem. The combined conclusions from both surveys allowed us to assess both species-specific and more general reactions of amphibian populations to multiple factors and hazards rampant in the urban environments of Israel. This work demonstrates how the unique advantages of citizen science can produce results with significant conservation implications, all while promoting conservation by connecting the community to its neighboring wildlife.

### 1.2 Study species

*Salamandra infraimmaculata* (Marten, 1885), or the near Eastern fire salamander, is a relatively large salamander species found in the near east, with Turkey as the northern border of its distribution and Israel as the southern (Degani et al., 2019; Steinfartz et al., 2000). This species is considered to be “Near Threatened” globally (Papenfuss et al., 2008), and endangered in Israel (Dolev & Perevolotsky, 2004). After mating, the females gestate for several months before spawning 10-200 larvea into the aquatic breeding site (Sharon et al., 1997). The juveniles usually spend several months in the aquatic phase of their lives, but the age and size at the time of the metamorphosis into the terrestrial adult phase vary significantly with environmental conditions (Goldberg et al., 2012). Most adult salamanders do not travel further than 10 meters during their foraging activity, though rare dispersal events of over 1 kilometer were documented (Bar-David et al., 2007). Adult salamanders can live up to 20 years but usually reach their nearly-maximal body-size by the 9^th^ or 10^th^ year of their lives (Warburg, 2007, 2008). Israeli *S. infraimmaculata* populations breeding in ephemeral and permanent water sources were found to vary significantly in their population size, with the former having typical population sizes of 30 to 130, and the latter sometimes numbering over 500 (Segev et al., 2010). An analysis performed on the distribution of salamanders in the Carmel found that their occupancy of urban and rural breeding sites is not significantly different (Blank & Blaustein, 2012).

*Bufotes variabilis* is a green toad species native to Israel (previously classified as *Bufotes viridis* or *Bufo viridis*)(Meiri et al., 2019). Toads of this species usually, but not exclusively, breed in ephemeral water bodies (Elron, 2007). The mating takes place in the water, during which the female may lay more than 8000 eggs (Elron, 2007). The tadpoles complete their metamorphosis within 3 months on average and reach sexual maturity at the age of two to three years. The mean lifespan is six years for males and four years for females(Elron, 2007). This toad species was once the most commonly sighted amphibian species in Israel (Bodenheimer, 1935; Nevo & Schneider, 1976), but its population declined significantly over the last decades (Dolev & Perevolotsky, 2004; Elron, 2007), and it is now considered to be locally endangered (Meiri et al., 2019).

## 2. Results

In this section we present the results of our analysis of two datasets. The first is the product of the Haifa Salamander Monitoring project, a citizen science program that aimed to monitor two *S. infraimmaculata* populations within the city of Haifa. The survey was first executed in 2016, in the Haifa educational zoo and the adjacent stream, Wadi Lotem. This population was previously monitored to some extent by the staff of the Haifa zoo and increasing concerns regarding its decline prompted a more thorough survey in the form of this citizen science program. This salamander population is thought to have originally been breeding in the small, ephemeral water bodies along the Wadi Lotem stream; another possibility is that it was introduced prior to the 1980’s from other sites in Israel via escapees from zoo captivity. Several additional potential breeding sites have existed at least since the 1980’s in the form of artificial ponds within the zoo. Some of these ponds contain invasive predatory fish, as well as dense vegetation and algae. It is feared that the exotic vegetation covering the water surface leads to anoxic conditions during certain times of the year, possibly rendering these ponds unsuitable for salamander breeding and even turning them into ecological traps. In the following year, 2017, the survey was extended to an additional site, a stream named Wadi Ahuza, three kilometers to the north of Haifa zoo. This site suffered a wildfire in 2016, and one of the project’s goals was to assess the wildfire’s impact on the resident salamander population.

The main component of the project was a terrestrial survey, which included two constant routes in Wadi Lotem/Haifa zoo and one constant route in Wadi Ahuza. All surveys took place following rainfall, and all salamanders that were spotted by the volunteers were measured, weighed, and photographed. The surveyors additionally inspected the potential breeding sites for the presence of both adult salamanders and salamander larvea during each survey night.

The source of our second dataset is a survey we conducted of two *B. variabilis* populations around the city of Jerusalem, following the same protocol as the salamander surveys, though without the assistance of volunteers. The focal site was the Gazelle Valley urban nature park within the city of Jerusalem. Like the Haifa zoo/wadi Lotem salamander population, the toads of Gazelle Valley had originally bred in the small, ephemeral water bodies along the two streams passing through the site. In 2015, an urban nature park was established in Gazelle Valley, and several artificial ponds, both ephemeral and permanent, were established. As in Haifa zoo, the presence of invasive predatory fish in these ponds, as well as a large population of the Levant water frog (*Pelophylax bedriagae*) which arrived at the site since the introduction of permanent water bodies, are feared to turn these ponds into potential ecological traps. As a control, we performed a survey following the same protocol at a site in which existed, according to our preliminary survey, a large population of *B. variabilis*: the Ein Hemmed national park, 8 kilometers northwest of Gazelle Valley. The presence/absence of adults and tadpoles of *B. variabilis* was documented in six additional sites around Jerusalem.

### 2.1 S. infraimmaculata

#### 2.1.1 Salamander citizen science project

Between February 2016 and February 2022, the salamander monitoring project’s volunteers performed 89 surveys, 28 in Wadi Ahuza and 61 in Wadi Lotem. Surveys were performed during rainfall or within a day of its occurrence. The volunteers captured altogether 212 visually distinct individuals, 162 in Wadi Ahuza and 50 in Wadi Lotem. The mean number of individual salamanders encountered per survey was 14.96 ±10.60 SD in Wadi Ahuza and 3.87±3.22 SD in Wadi Lotem, with the difference in frequencies being statistically significant for α = 0.05 (Welch Two Sample t-test: t = -5.41, df = 29.65, p << 0.001).

The mean length of salamanders captured in the Lotem population (M = 24.98, SD = 3.44) was significantly larger than the length (M = 22.36, SD = 3.25) of salamanders captured in the Ahuza population (Welch Two Sample t-test: t = -4.4924, df = 63.299, p-value = 3.055e-05). The lengths of males (M=23.24, SD=2.50) and females (M=23.72, SD=2.21) did not differ significantly (Welch Two Sample t-test: t = -1.3645, df = 156.01, p-value = 0.1744). The proportion of sighted females in the Lotem population (0.26, n = 47) and in the Ahuza population (0.36, n = 152) did not differ significantly (Fisher’s Exact Test for Count Data: p-value = 0.2238).

In Wadi Ahuza, salamander tadpoles were sighted in both the ephemeral and permanent water bodies along the stream during all survey years. In Wadi Lotem / Haifa zoo, during the sampling season of 2016-2021, tadpoles were only found in one water body, a small artificial pond within the zoo.

#### 2.1.2 Movement and activity patterns

To understand the salamanders’ habitat use and site fidelity, and to identify possible dispersal events, we explored the salamander’s movement patterns by calculating the distances between all capture points for 68 *S. infraimmaculata* individuals for which the exact capture location was specified in more than one capture.

Of the individuals for which more than two exact locations were documented, 85% were found in the exact same location (within 1-20 meters) on a least two different occasions, 54% were found in the same location three times, and 19% were found in the same location more than five times. 37% of these individuals were found in the same exact location in more than one winter. All but five salamanders were captured while in terrestrial activity. 64.8% of documented capture locations were within 100 meters of a known breeding site.

Additionally, we calculated the maximal displacement for each of these individuals using the maximal distance between two of its capture locations (see figure 1). We conducted a two-way ANOVA to test the effect of sex and population of origin on an individual salamander’s maximal documented displacement, with no statistically significant effects found for population (F(1) =0.12, p =0.73) and sex (F(1) =0.24, p =0.62) and no significant interaction found between their effects (F(1) =0.01, p =0.93).

**Figure 1.**
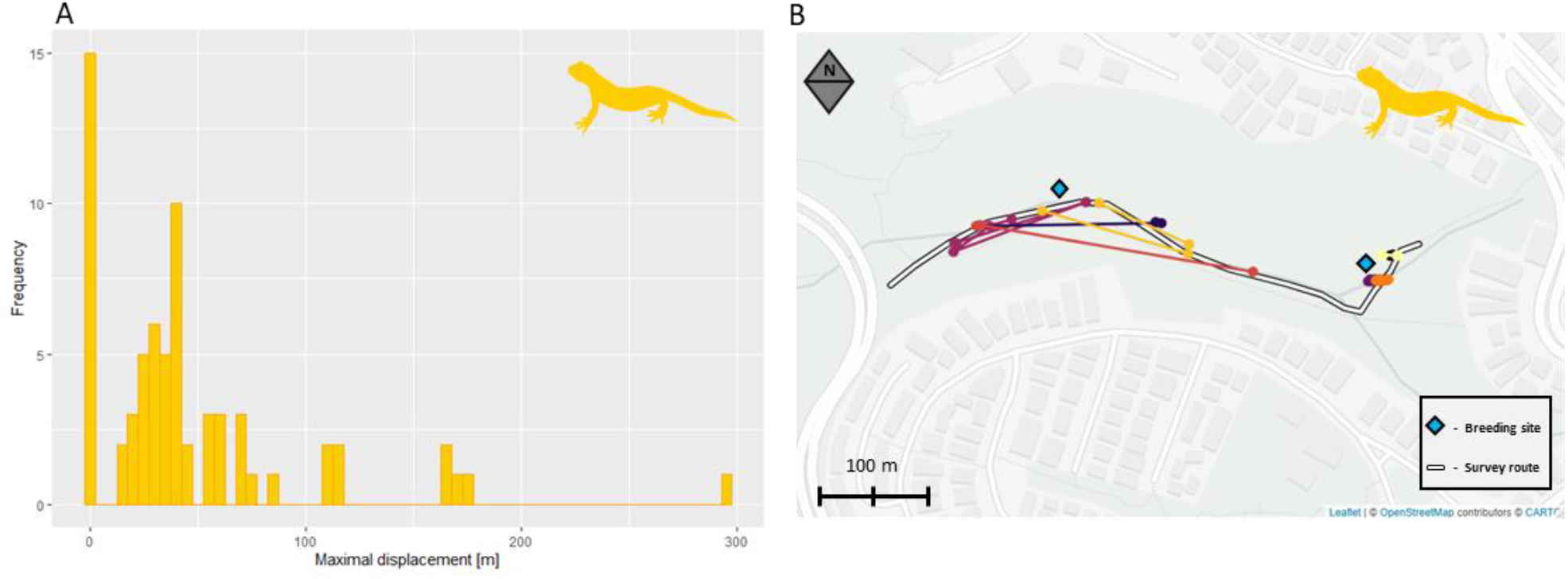
*S. infraimmaculata* movement patterns-. A: A histogram of maximal documented displacement [m] of individuals with two or more than documented exact locations (n=68). B: Exact capture locations of 12 individuals from the Wadi Ahuza population. Points of the same color mark repeated captures of the same individual. Consecutive capture points of each individual are connected by lines. The individuals displayed were those with the four highest and four lowest maximal documented displacement values in the Wadi Ahuza population. Some points have been slightly shifted to enhance their visibility.

#### 2.1.3 Growth rate and age estimation

The age structure of a population can be crucial to our understating of its viability. Previous studies have described the growth of tadpoles or juveniles of this species and adults of other species in the same genus as an exponential function (Najbar et al., 2020; Warburg et al., 1979). However, no reliable method for estimating the age of adult salamanders of the species *S. infraimmaculata* is currently available. We derived a rough estimate based on our own data. The growth rate of *S. infraimmaculata* is nonlinear, and we therefore used the repeated measurements of 84 individual salamanders over multiple years to quantify the link between their length and their estimated growth rate. We used it to develop a formula for estimating the age of *S. infraimmaculata* individuals using their total length.

Similarly to Warburg (2008) we performed a linear regression (see figure 2A) between the salamanders’ growth rate (*[length at year x+1]-[length at year x]*) and length from snout to tail tip (*([length at year x+1] + [length at year x])/2*) on all individuals which were measured in two consecutive years. For individuals that were measured in 2 consecutive seasons more than once, we used the earliest measurement, to account for the scarcity of measurements of shorter (and therefore, younger) individuals in the data. Measurements of two individuals indicated that they shrunk by 2.5 cm between sampling seasons. Based on existing knowledge of the species (Warburg, 2008) these were marked as errors that were likely to be caused by tail truncation due to injurie, and therefore excluded from further analysis.

**Figure 2.**
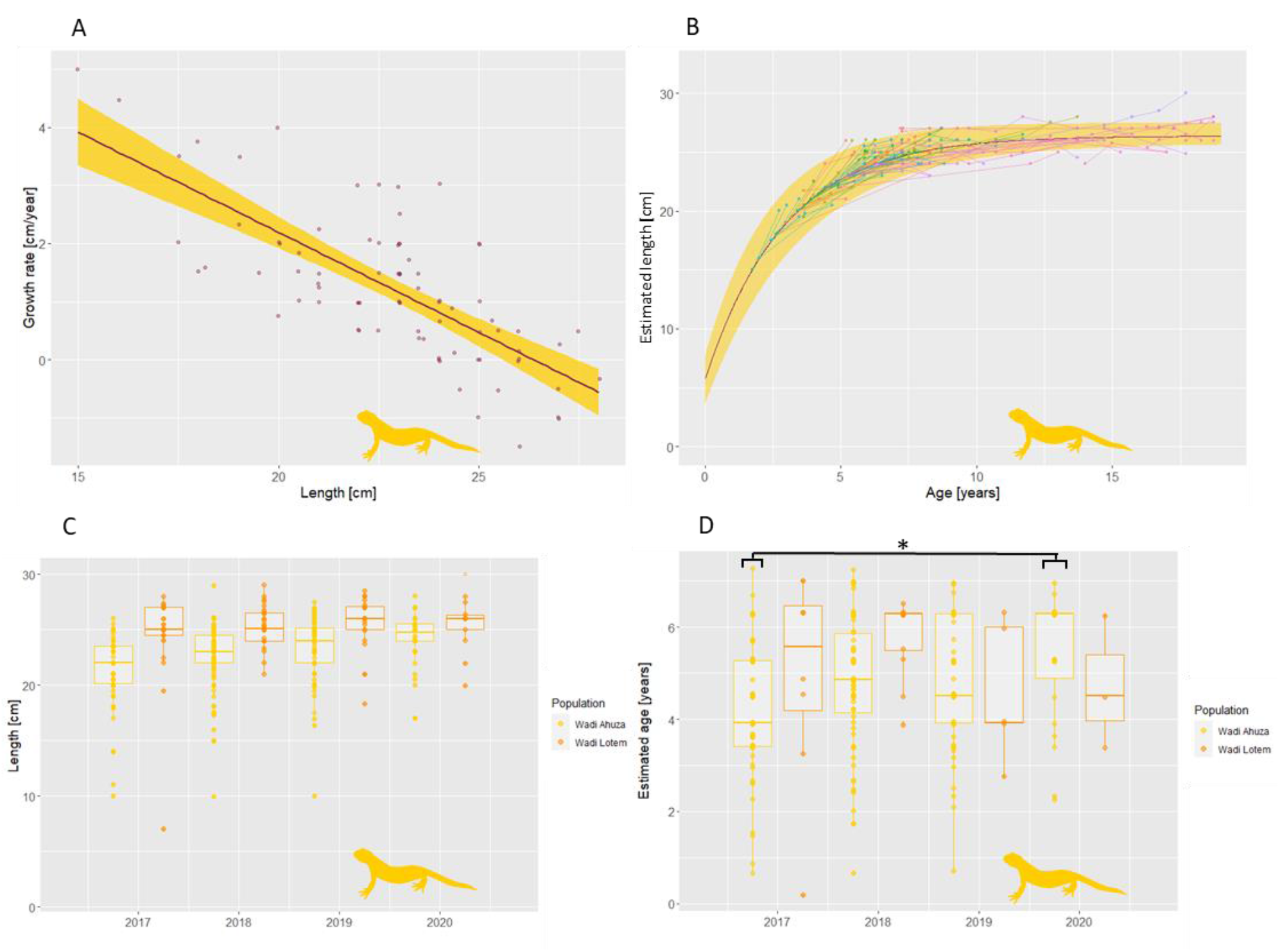
*S. infraimmaculata* growth rate and aging A: Individual salamanders’ mean length over two consecutive years (x-axis) and the difference in mean length between the two years (y-axis). The points show the raw data for all salamanders that were measured in two consecutive years (n = 68). The trend line and area around the curve represent the fitted linear regression (Growth rate∼ Length) and its 95% CI. B: the growth function L(t) which was derived from the linear regression. The curve represents the estimated mean length as a function of the salamander’s age and the area around the curve represents a bootstrap 95% CI for that mean. The colored points show the data for all salamanders that were measured more than once (n = 106), with each salamander’s initial age estimated by its length upon its first measurement. The ages of salamanders that were longer than the upper limit we set for age estimation (24.92 cm) were adjusted to show the likely age distribution in the population more accurately, using the mean adult mortality /year estimated in our capture-recapture analysis (0.177) (see text). Points of the same color mark repeated measurements of the same individual. The consecutive points of each individual are connected by lines. C: Scatterplot of the lengths of all measured individuals in the two studied salamander populations in the years 2017-2020. D: Scatterplot of the estimated ages of all measured salamanders that were short enough for age estimation (length < 24.92 cm). The mean age of the individuals in the Wadi Ahuza population in 2017 was significantly lower than the age of the individuals of the same population in 2020 (Tukey’s honest significance test, see text).

The fitted regression model was:

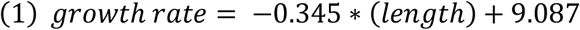

If we mark the length of a salamander at the age *t* as *L*_(*t*)_, we can express the linear model as:

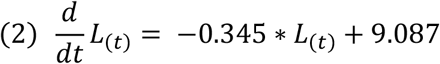

The overall regression was statistically significant (R^2^ =0.44, F (1,82) =66.9, p = 3.081e-12, see Fig. 2A). we used the estimated slope and intercept to solve equation 2 (see Methods) and acquire the growth function, *L*_(*t*)_, with the growth coefficient (k) set at -0.345 and the asymptotic length (A) set at 26.373, using the length upon completion of metamorphosis from Goldberg et al. (2012) as *L*_(*0*)_:

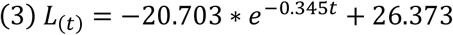

We used permutation tests to determine if these parameters’ estimations differed significantly between sexes and populations. Between males and females, the differences in both the growth coefficient (10,000 iterations, Δk =0.028, p = 0.77) and the asymptotic lengths (10,000 iterations, ΔA = 0.97, p = 0.38) were non-significant. Between the Ahuza and Lotem populations the differences in both the growth coefficient (10,000 iterations, Δk =0.060, p = 0.51) and the asymptotic lengths (10,000 iterations, ΔA = 0.336, p = 0.74) were non-significant as well.

The 95% CIs for parameters estimated from the data were estimated using bootstrap and were corrected for bias (10,000 iterations, k :(estimate = -0.345, upper = -0.268, lower = -0.414), A: (estimate = 26.373, upper = 27.433, lower =25.645)). The CI for the starting length was derived by simulating a normal distribution using a mean (=5.67) and SD (=1.04) from existing literature (Goldberg et al., 2012).

We then used the reversed function for *L*_(*t*)_,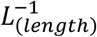, as an estimate of a salamander’s age based on its total length. However, the function *L*_(*t*)_ approaches an asymptote, from a certain point 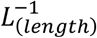 becomes extremely sensitive to small differences in length, producing unreliable results. We set the upper limit of detectable growth, based on the resolution of the length measurements in the survey, as 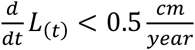. We, therefore, determined the upper limit of estimable ages, using the derived parameters, to be 7.35 years, with the respective length being 24.92 cm. Salamanders longer than that were given the age of 7.35 years but excluded from further analysis regarding the age composition of the two populations since they are irrelevant for comparing the recruitment of juveniles during the survey years. We performed a two-way ANOVA on the estimated ages with respect to both the population and survey year, both the population’s effect (p<0.01) and the survey year’s effect(p<0.01) were found to be significant, though their interaction was not (p = 0.143). We then performed Tukey’s HSD test, and a significant difference was found within the Ahuza population, with the mean age in the year 2020 higher than in the year 2017 (difference =1.27, adjusted p-value =0.02). Three other significant differences were found between Ahuza and Lotem subgroups.

#### 2.1.4 Capture-Recapture analysis

To estimate demographic parameters such as survival rate and population size, we performed a capture-recapture analysis using the “robust design – Huggins *p* and *c*” model structure in the program MARK (White & Burnham, 1999).

We constructed 28 a priori capture-recapture population models, with different combinations of length, sex, air temperature, relative humidity, sampling season, and sampling occasion as predictor variables. Of these models, the data conclusively supported (total AICc weight > 0.99) models that included an effect of length on both capture probability (ß = 0.104449879, 95% CI = 0.060749107 (lower) - 0.148150723 (upper), LOGIT link function parameter) and survivorship between consecutive years (ß = 0.0565, 95% CI = 0.040207031 (lower) - 0.072509265 (upper), LOGIT link function parameter), and both of those effects were found to be statically significant (p _ß=0_ < 0.05) in all of the supported models. The effects of air temperature and relative humidity were both included in all of the supported models and were both found to be statistically significant (p _ß=0_ < 0.05). The average survival rate between consecutive years weighted by each model’s AICc weight was 0.775 (95% CI = 0.699 (lower) - 0.837(upper)) for the Wadi Ahuza population and 0.766 (95% CI = 0.683 (lower) - 0.832(upper)) for the Wadi Lotem population. The estimated population size for each sampling season did not change significantly over time in either population (figure 3A). Though females were underrepresented (19% of observations were females, 70% males, and the rest were of unknown sex or juvenile). all models containing an effect of sex on capture probability failed to produce a ß_sex_ parameter estimate, possibly due to the sample size for females being too small. These models were therefore excluded from the analysis, but this bias (Which possibly stems from inherent bias in the capture method cause by the males’ behavior during mating season, for example : Hall, 1977; Marsh & Goicochea, 2003; Marvin, 1996; Salvidio, 2008) may have resulted in an underestimation of the population size by our analysis. If we were to assume that this underestimation is proportional to the sex bias among our captured individuals, the corrected population sizes would larger by approximately 50%. The population size was additionally estimated based on the non-parametric approach described in (Chao et al., 1992). See all population size estimates in table S1.

**Figure 3:**
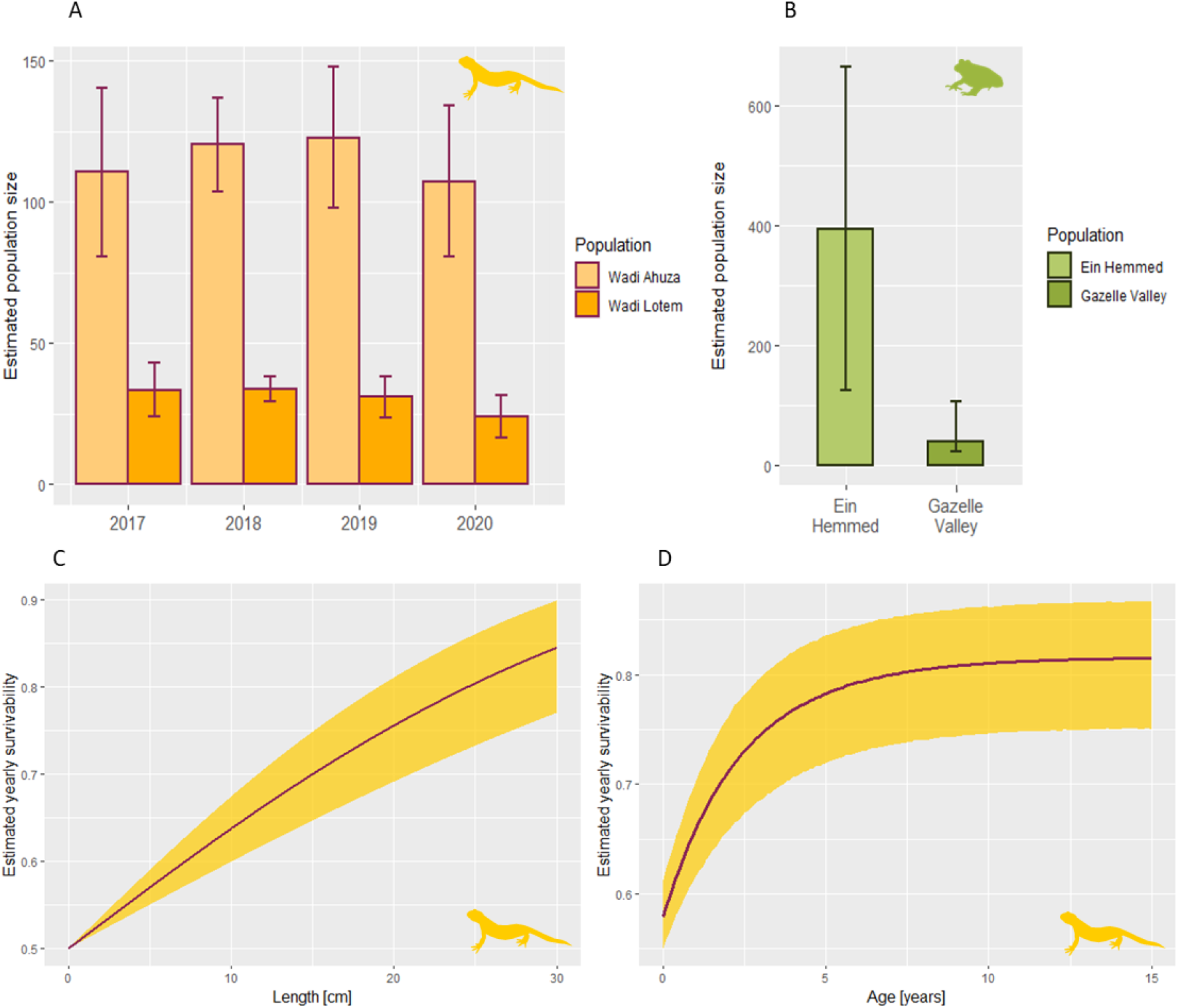
Parameters estimated using Capture-Recapture analysis. Estimated sizes of the two studied S. *infraimmaculata* populations from 2017 to 2020 (A) and of the two studied *B. variabilis* populations in 2020 (B). The error bars represent 95% CIs. C: the link function between Length and the probability of survival between consecutive years estimated using the Capture-Recapture analysis (LOGIT (−0.0565 * L) = S), the area around the curve represents 95% CI. D: Survival rates of salamanders along their lifetime simulated (10,000 iterations) using the derive age-to-length and length-to-survival rate function parameters and their respective CIs. The area around the curve represents the range between the 5% and 95% quantiles of the simulation results.

#### 2.1.5 The effect of age on the survival rate of *S. infraimmaculata*

Using the estimate and the CI for the effect of length on the survival rate of individual salamanders we derived from the Capture-Recapture analysis (ß = 0.0565) we plotted the link between length and survival rate (figure 3C). We then simulated the survival rate for each age group by randomly sampling values from within the 95% confidence intervals of the estimated age-to-length and length-to-survival rate function parameters (10,000 iterations, see figure 3A).

### 2.2 B. variabilis

#### 2.2.1 Toad survey

We conducted 21 nocturnal surveys overall in search of B. variabilis, twelve in Gazelle Valley urban nature park and nine in Ein Hemmed national park, between August 2020 and May 2021. In Gazelle Valley and Ein Hemmed, we captured 18 and 79 individuals respectively, and all individuals had distinct enough patterns to allow identification upon recapture except three individuals captured in Ein Hemmed. Over 76% of the individuals captured in Ein Hemmed were males caught in or nearby a water body, displaying characteristic courting behaviors (floating at the water’s surface, emitting mating calls). No individuals captured in Gazelle Valley were inside a water body. Two individuals emitting mating calls inside a water body were observed during the survey but not captured. *B. variabilis* tadpoles were observed in Ein Hemmed in three surveys, in multiple water bodies inside the park, numbering in the thousands on each occasion. In Gazelle Valley, only one group of 435 *B. variabilis* tadpoles was observed on one occasion (manually counted from a photograph), but no tadpoles or metamorphs were observed in the same water body in subsequent surveys.

We observed 16 metamorphs in Ein Hemmed. The discrepancy between the many thousands of tadpoles that were observed and the relatively small number of metamorphs can be possibly accounted for by the metamorphs’ tendency to spend most of their time in hiding in the weeks following the metamorphosis (Elron, 2007). No metamorphs were observed in Gazelle Valley. We could not detect metamorph recaptures due to the lack of a reliably stable and distinct dorsal pattern at this age. The mean number of individual toads encountered per hour of surveying was 10.56 ±9.71 SD in Ein Hemmed and 1.69 ±2.96 SD in Gazelle Valley, with the difference in frequencies being statistically significant for α = 0.05 (Two-Tailed t-test Two-Sample Assuming Unequal Variances: t = 2.50, df = 8, p = 0.037).

#### 2.2.2 Capture-Recapture analysis

From the six models developed for the Ein Hemmed population, the two models supported by the data (total AICc weight > 0.99) included an effect of survey duration on capture probability, which was statically significant in both models (p _ß=0_ < 0.05). The less supported of these two models (AICc weight = 0.267) included an effect of sex on capture probability, but this effect was not statistically significant (p _ß=0_ > 0.05). The average estimate for the Ein Hemmed population size, weighted by the AICc, was 395 (95% CI = 125 (lower) - 666(upper)) (figure 3B). Female captures were rare in the Ein Hemmed survey (14% of observations), and the estimated parameter for the effect of sex on capture probability was too small to affect the population size estimation, leading to a possibility of an underestimation of the population size by our model. If we were to assume that this underestimation is proportional to the sex bias in our captures, the corrected population size for Ein Hemmed would be approximately 50% larger (∼600 individuals).

For the Gazelle Valley population, the estimated population size was 40 (95% CI = 23 (lower) – 106 (upper)) (figure 3B). Due to the small sample size (21 total observations, 3 individuals captured more than once) we did not analyze the effect of environmental and individual covariates on capture probability in this population.

## 3. Discussion

In this section, we discuss the conclusions from our analysis, and their possible Implications for the conservation of amphibian species in urban environments and outside of them. We will further discuss the promise our results show for exploring phenomena and dynamics that were previously out of reach using intensive, non-invasive citizen science surveys. Finally, we will offer an outline for future research programs, aimed at informing conservation efforts by harnessing the unique advantages of citizen science.

A useful result of this capture-recapture analysis is our estimate of the studied populations’ sizes. While the sizes of both populations were stable throughout the project’s duration, the salamander population in Wadi Lotem and the adjacent Haifa Zoo was consistently smaller than the population of Wadi Ahuza. The population size of Wadi Lotem is similar in size to *S. infraimmaculata* populations in the Carmel which breed in ephemeral ponds, despite the presence of permanent water bodies inside Haifa Zoo, opposite to the trend described in Segev et al. (2010). A possible conclusion is that some attributes of these water bodies, most likely the continued presence of fish, prevents them from functioning as breeding sites for the local salamanders.

Many of our results agreed with previous studies on the salamanders of Carmel, including the scale of our estimated population sizes, the maximal documented displacements, site fidelity, and the relationship between age and length (Bar-David et al., 2007; Segev et al., 2010; Sinai et al., 2020; Warburg, 2008). This agreement is a possible indication that the results achieved by this citizen science project are similar in their reliability and precision to those acquired by the more traditional surveys conducted by researchers while adding the educational and societal advantages offered by citizen science.

Warburg (2008) studied *S. infraimmaculata* in the Carmel for decades, using the link between length and growth rate, amongst other measures, to study the growth patterns of these salamanders and the possibility of length-based age estimation. While utilizing a similar methodology, we managed to replicate Warburg’s results regarding the changes in the growth rates of these salamanders over the course of their lives, namely, that the majority of these salamanders’ growth occurs in their first eight years of life. Warburg, however, determined that the random variance in the salamanders’ lengths does not allow for reliable length-based age estimation in this species. Similarly, we avoided drawing conclusions from our length-based age estimation regarding specific individuals, or from the differences in length between the two studied populations. However, we assert that a change in the mean length-based age estimate in a specific population between consecutive years, such as the significant change we found in the Wadi Ahuza population, is a reliable measure for changes in recruitment, and therefore an informative tool for research and conservation. Used cautiously, the age estimate produced by our formula can be informative on its own, at least for populations of a similar genetic makeup and habitat, such as other salamander population in the Carmel.

We quantified the underrepresentation of juvenile salamanders in our survey, a result which may be applicable in future studies, improving our approximations of the age structures of *S. infraimmaculata* populations. This underrepresentation might have made it difficult to detect the rate of juvenile recruitment. Nevertheless, the sizes of both populations were stable during this period, while the estimated mortality was over 10%, which might mean recruitment is occurring to some extent, but can also indicate an error in our assessment.

Furthermore, our analysis shows that salamanders of this species do not reach their peak survivorship until they complete the majority of their growth, around the age of eight years. relatively low survivorship during their first years of life might mean that a loss of adult salamanders cannot be compensated for by juvenile recruitment without several consecutive years of successful breeding and terrestrial habitats stable enough to allow a sufficient fraction of the sensitive juvenile salamanders to complete their growth. A possible major implication for future conservation efforts is that adult, fully grown salamanders should be thought of as a vital resource for sustaining the population, that should be prioritized alongside the breeding sites.

Like the two salamander populations, the toad populations differed substantially in their sizes; in both cases, the smaller population was found at a location in which the aquatic breeding sites were repeatedly disturbed in the years before the survey and had a consistent presence of predatory fish. Unlike the smaller populations, both larger populations of each species had access to multiple small, stable, and diverse breeding sites. The Ein Hemmed toad population was both the largest amphibian population in this study and the only population that showed clear signs of successful breeding. Unsurprisingly, Ein Hemmed is the most distant site from urban centers and the only site located within a protected area. Gazelle valley was the site with the most diverse amphibian community in this study, Yet the most threatened species in it, which was the focus of our survey, had a small population that showed no signs of successful juvenile recruitment during the survey season.

As we repeatedly emphasized in this work, the breeding site’s characteristics prove to be critical; a major concern regarding both Gazelle Valley and Wadi Lotem is that the fish-containing, non-ephemeral ponds function as ecological traps. One of the unique features of amphibians in the semi-arid levant is the ability of amphibians to breed in small, ephemeral aquatic habitats, and the presence of permanent ponds is not necessarily beneficial and can even be detrimental. The presence of fish, and particularly of the invasive *Gambusia affinis*, which was shown to prey on tadpoles of both *S. infraimmaculata* and *B. variabilis* in laboratory experiments (Elron, 2007; Segev et al., 2009), might be the determining factor. The implication arising from these aspects of our study is that conservationists in this region should consider providing the target populations with multiple small, diverse breeding sites, to minimize the chances that a single large unsuitable breeding site will doom the entire population.

One of the main challenges in identifying threats to amphibian populations is the difficulty to trap enough individuals multiple times, which is necessary for almost all estimations of movement patterns, population dynamics, and life history traits. The Haifa salamander monitoring project managed to address this with a simple format. Here, a small group of trained researchers and a larger team of volunteer trackers formed an effective surveying unit that sampled its research sites consistently, thoroughly, and over a long period of time. A common criticism of surveys based on citizen science is that their resolution is often limited to presence/absence and species richness data. The approach developed in the Haifa salamander monitoring project, however, managed to provide details on the population structure and life history, which are paramount to making informed management decisions regarding these populations and their environments. Together with our green toads’ survey, our analysis highlights that this may be crucial, because species richness is not a sufficient measure of a site’s suitability for sustaining amphibian populations.

Finally, we suggest an outline for future monitoring of urban amphibians, based on these projects and the conclusions stemming from them. Physically Handling any wild animal has a potential of causing it distress and even lasting damage. In the toad survey, both recognition of recaptures and length measurements were achieved using pictures alone. We therefore assert that there may be no need for capturing and handling animals for such capture-mark-recapture surveys. A possible promising venue for future citizen science programs may be to outline specific survey routes at multiple sites and allow volunteers to survey the sites that are in their immediate vicinity independently. The volunteers can be instructed in how to photograph the amphibians alongside standard scale cards while causing minimal interference in the animals’ activities, allowing them to then survey the adults and tadpoles along specific routes on a regular basis. By continuously recruiting volunteers to regularly survey many sites, it may be possible to gather the crucial data conservationists will require to effectively protect the amphibians of the Mediterranean.

## Methods

### *Salamandra infraimmaculata* Citizen science project

The Haifa Salamander Monitoring Project was initiated and led by local volunteers, collaborating with several local organizations dealing with conservation, research, and environmentalism. An initial group of volunteers with research and conservation backgrounds designed the survey protocol and routes and recruited a larger team of volunteers, trained them to track and safely capture the salamanders, and supervised the teams performing the surveys.

The volunteers walked in teams of 3-4 along a constant route in each survey site, always at nighttime, scanned the path using flashlights, and documented every salamander they spotted along their path. They measured its length (snout to tip of tail, and occasionally snout-to-vent), weighed it, photographed its unique dorsal spot pattern for later individual identification, and documented the capture location. All measurements were performed by a specific team member with the required training. The volunteers additionally documented the temperature and relative humidity during each survey. The data includes the observations from the main survey (Wadi Lotem: 2016-2022, Wadi Ahuza: 2017-2021), casual observations of salamanders in the city of Haifa during the same years, and observations of salamanders captured from the Wadi Lotem population by members of the Haifa Zoo staff during the years 2007-2009. The volunteers additionally noted possible hazards to the salamanders spotted during the survey, as well as the presence of salamander larvea or other amphibian species in the breeding sites.

In addition to the engagement of the survey participants, the project included additional efforts to inform and engage the community, including posting signs in the breeding sites, free lectures about the salamanders’ biology, social media activity, and the operation of information booths around the city of Haifa. The surveys were conducted under INPA permits number 2018/42061, 2019/42368, and 2020/42648.

### Study site

The *S. infraimmaculata* survey took place in two sites: Wadi Lotem stream/ Haifa Zoo between 2016 and 2022 and Wadi Ahuza stream between 2017 and 2022. Both sites are directly adjacent to the city of Haifa, Israel. This area, within the Carmel Mountain range of northern Israel, is the southernmost edge the global distribution of *S. infraimmaculata*. The Carmel was originally covered by Mediterranean woodland, with numerous caverns and springs as suitable habitats, shelters, and breeding sites for salamanders of this species. The urbanization of the region which occurred gradually since the mid-20^th^ century altered and disturbed many of these habitats, and its effects on the salamander population are not fully understood (Degani et al., 2019). The habitats in the study sites were both altered by their proximity to the city of Haifa, including wildfire, water pollution, water body alterations, discarded waste, and the presence of feral cats and dogs. Concerns regarding the state of the resident salamander populations motivated the Haifa Salamander Monitoring Project.

### Data analysis

In both surveys, we manually recognized recaptures of individuals of the species *B. variabilis* and *S. infraimmaculata* from the pictures taken during the surveys using their unique dorsal spot patterns. The resulting datasets included a list of all documented captures, with the location, time, and all measures taken from each individual, as well as a list of all surveys performed, with each survey’s time, location, number of individuals captured, and all other available details.

The data from the last salamander sampling season in Wadi Ahuza (winter of 2021-2022) was not yet processed at the time of this analysis and was therefore excluded from it. To test simple effects of the sex or population, we averaged multiple measurements of the same individuals (unless otherwise stated). The data was analyzed in the Rstudio platform (Allaire, 2012) using the packages toolsForAtlas, ggplot2 (Wickham et al., 2016), Dplyr (Mailund, 2019),tidyverse (Wickham & Wickham, 2017) ggpubr (Kassambara & Kassambara, 2020), Boot (Canty, 2002), leaflet (Cheng et al., 2019), and MASS (Ripley et al., 2013).

### Growth rate and age estimation

To determine the salamanders’ ages, we sought to derive a function that describes their growth throughout their lives (will be marked as L_(t)_). In this analysis, we could not use direct regression methods since we do not have any data about any of the salamanders’ ages. However, the L_(t)_ function derived for similar species (Najbar et al., 2020) and for larvea of salamanders of this species (Warburg et al., 1979) were exponential. Therefore, if we assume the L_(t)_ function is exponential, we can derive this differential equation:

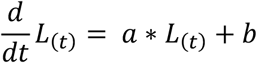

which describes a linear link between the salamander’s length at time *t* and its growth rate at time *t*. Since we had multiple observations of salamanders from consecutive years, a linear regression between growth rate (*[length at year x+1]-[length at year x]*) and length (*([length at year x+1] + [length at year x])/2*) could be performed to derive *a* and *b*.

The solution for the differential equation is:

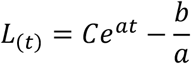

Where *a* is the growth constant (k), 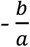 is the asymptotic length (A) and C is the difference between the asymptotic length and the initial length, which was taken from previous studies (Goldberg et al., 2012). From the same parameters, a reverse growth function-length to age – could be derived:

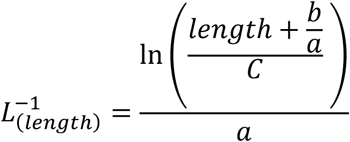

and used to convert the salamanders’ measured lengths to age estimations. However, since *L*_(*t*)_ approaches an asymptote, from a certain point 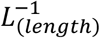 becomes extremely sensitive to small differences in length, producing unreliable results. We determined the upper limits of the reliability of 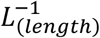 as the point at which the yearly growth rate 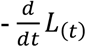 - is 0.5 cm/year since 0.5 cm was the measurement resolution in the survey.

### Capture-Recapture analysis

For the Capture-Recapture analysis, we excluded all observations that did not occur during the standard-effort surveys, and all individuals that did not have a distinct enough pattern for identification upon recapture. For surveys in which air temperature, relative humidity, or survey duration was missing, the average for all other surveys was used. Since our planned model structure for the *S. infraimmaculata* dataset required the individual covariates (sex, length) will be listed for each sampling season for all individuals, and not all individuals were captured and measured in all seasons, the missing lengths were estimated using the growth function we derived from the data.

All datasets were analyzed using the program MARK (White & Burnham, 1999). For the *S. infraimmaculata* data, we constructed 28 a priori population models, with different combinations of length, sex (individual covariates), air temperature, relative humidity (environmental covariates), sampling season, and sampling occasion (time dependency) as predictor variables the probabilities for individual being captured in a specific occasion, surviving from one sampling season to the next, immigrating between sampling seasons, and emigrating between sampling seasons. The models were implemented using the “robust design – Huggins *p* and *c*” (under the assumption that captures do not alter the salamander’s behavior in a way that affects future captures: *p*= *c*) model structure available in MARK.

To display individuals that were longer than the maximal length allowing age estimation (24.92 cm) on the growth curve in fig. 2B, we used the estimated mean survival rate for fully-grown adults (0.823) to simulate the age distribution of the subpopulation of salamanders longer than 24.92 cm (under the assumption that the recruitment is constant for this subpopulation) and used this distribution to produce a rough estimation of their age, based on the rank order of their lengths.

### *Bufotes variabilis* survey

The study sites of the *B. variabilis* survey were the Gazelle Valley Urban Nature Park inside the city of Jerusalem and the Ein Hemmed National park in the adjacent Judean Hills region. Gazelle Valley was a highly disturbed habitat for several years, with large amounts of waste inside and outside of the water bodies. The urban nature park on the site was established in 2015, and vast rehabilitation efforts were devoted to the aquatic habitats, including a water suspension and circulation system which added permanent ponds of differing sizes to the preexisting small ephemeral water bodies. However, it is possible that the ambient pollution present in the urban runoff water which flow into those water bodies, as well as the presence of the invasive predatory fish species *Gambusia affinis*, does not allow some amphibian species to breed in them, to a degree they may constitute ecological traps. Our survey sought to assess the influence of these changes on the preexisting amphibian population, by determining the population size and composition and comparing them to those of a control population. The chosen control population was the toad population of Ein Hemmed National park, located near a suburban area of the Judean Hills, in which a similar water suspension and circulation system have existed for over a decade, and in which large numbers of *B. variabilis* were observed in our preliminary survey.

### Field survey

We performed a series of nighttime surveys in Gazelle Valley urban nature park and in Ein Hemmed national park in order to assess the population size and composition of *B. variabilis* in both sites, while documenting their breeding activity, possible hazards, and the presence of other amphibian species. We recorded all amphibian observations and their location but only members of the species *Bufotes variabilis* and *Hyla savignyi* were photographed alongside a scale bar, and their snout-to-vent lengths were measured using the scaled photographs. We included in the route the approachable shorelines of all known water bodies in both sites and recorded the presence of tadpoles and eggs, as well as breeding activity by adults of the surveyed species. We performed the surveys along a constant route, during the evening hours, and documented the air temperature, relative humidity, survey start time, and duration for each survey. No more than two people took part in each survey.

We additionally performed two daytime surveys in the water bodies of Gazelle Valley Urban Nature Park and recorded the amounts of amphibian tadpoles, eggs, and other marine organisms acquired after 10 net swipes, in three different depth categories.

For the *B. variabilis* data from Ein Hemmed National park, we constructed 6 a priori models with combinations of survey duration, sampling occasion, sex, and length as predictor variables of the probability of an individual being captured on a specific occasion. The models were implemented using the “Closed capture - Huggins *p* and *c*” (also assuming *p*= *c*) available in MARK. The *B. variabilis* data from Gazelle Valley Urban Nature Park was analyzed using the “Closed capture - full-likelihood” model structure available in MARK.

## Author contributions

O.D. carried out the *Bufotes variabilis* surveys, conducted the analysis, and wrote the manuscript. O.K and D.O. planned the study. O.K. supervised the research and commented on the manuscript. O.R, A.B.L, G.K and T.K.P planned and executed the *Salamandra infraimmaculata* surveys and commented on the manuscript. N.L.A and R.V accompanied the *Salamandra infraimmaculata* surveys on behalf of the SPNI and commented on the manuscript.

## Acknowledgements

We first and foremost thank the Haifa Salamander survey volunteers who, with great dedication, carried out the salamander surveys in the coldest and rainiest nights of winter.

We thank Liran Sagi, for his valuable guidance and assistance with the MARK analysis, All members of the Kolodny lab for their help and input, and Assaf Shwartz, Avi Bar Massada, Rachel (Ekly) Ben Shlomo, and Amir Balaban for extensive discussions that inspired this study.

We thank the Israel Nature and Parks Authority, Society for Protection of Nature in Israel, ”Yarok Ba’Lev” NGO, the Haifa Educational Zoo, the Municipality of Haifa, and Gazelle Valley Urban Nature Park for their essential cooperation and assistance in carrying out the surveys and the management of the data from the Haifa Salamander survey, and to the Haifa community of the SPNI in particular for their help and support of the Haifa Salamander project over the years. We thank Yael Hammerman, Avishai Shoresh, Noemi Cohen Kappach, Chen Chaim, for their help and support in carrying out the toad surveys, analyzing preliminary datasets, and involvement in various phases of the citizen science project.

## Conflict of Interest (required for all article types)

The authors have no conflict of interest to disclose.

## Data availability statement

All data is available from the corresponding author by request.

